# Weighted Shoes in the Wild: Initial Insights into the Relationship Between the Effort of Walking and the Amount of Walking Performed

**DOI:** 10.1101/2021.11.18.469130

**Authors:** Mailing R. Wu, Peter G. Adamczyk, Steven H. Collins

## Abstract

**Purpose:** Walking comprises a large portion of active energy expenditure in humans. Interventions that increase or decrease the energy expended on each step, such as ankle weights or energy-saving orthoses, may therefore strongly impact fitness. The overall effect of increasing or decreasing per-step energy use is unclear, however, because people may choose to walk less or more, respectively, in response.

**Methods:** In this study, healthy college students with normal body mass index wore weighted and unweighted shoes embedded with an inertial measurement unit for one week each. Community-based walking data were analyzed for number of steps, distance traveled and walking speed. Oxygen consumption using each set of shoes at a range of speeds were measured in a laboratory setting and used to estimate metabolic energy expended during community-based walking. A survey measured subjective response to each pair of shoes.

**Results:** The weighted shoes increased per-step energy cost by about 26%. Subjects strongly disliked the weighted shoes (P = 0.001) and found them tiring (P = 0.003). Despite this dislike, subjects did not significantly reduce distance walked (P = 0.6), number of steps (P = 0.7), or average speed (P = 0.9) compared to normal shoes. This led to a small but not statistically significant increase in energy expended during walking over a five-day period (12.3 ± 9.6% increase, P = 0.2). On the final collection day this trend appeared to reverse, with fewer steps taken and lower metabolic energy expended with the weighted shoes. Twenty-four subjects were recruited but only ten completed the protocol, with dislike of the weighted shoe condition being the primary reason for dropout.

**Conclusions:** Increasing the energy cost of each step led to greater energy expended through walking. However, there are indications that behavioral changes would be greater with a longer intervention or increased retention. For example, the large dropout rate suggests that some subjects avoided walking with the weighted shoes entirely, simply by leaving the study. Follow-on studies among patient groups may reveal a fitness benefit to either increasing or reducing the energy cost of walking.

## INTRODUCTION

The United States has recently seen a dramatic increase in the prevalence of obesity, with 75% of adults overweight and 41% having obesity.^1^ Overweight individuals and individuals with obesity have a high prevalence of comorbidities, particularly diabetes and hypertension.^2^ Complications associated with obesity have an estimated cost of $ 209.7 billion within the US healthcare system.^3^ Multiple studies have explored methods for lifestyle changes^4-7^ and adherence to exercise plans^8,9^ in an attempt to slow or reverse the obesity epidemic. People with obesity, and others with disabilities that affect the lower limbs, expend more energy per step, take fewer steps, and have lower physical activity levels than their normal body-mass-index or able-bodied counterparts, respectively.^10,11^

Humans spend a large portion of time in activities of moderate intensity, such as walking, across all age groups.^6^ Indeed, walking has been shown to be an effective method for promoting activity among sedentary groups, and people who walk more tend to have lower body mass indexes.^12^ People also adhere to walking routines more than other exercise regimens.^7,13^

In this study, we explored whether a modest increase in energy cost per step affected overall energy consumption from walking over a five day period compared to normal walking energy expenditure. While increasing the energy cost per step would increase net metabolic energy use if walking distance and speed were kept constant, overall energy cost may instead be reduced if the subject reduces overall activity due to the higher cost. We hypothesized that even a small increase in energy cost per step would cause the amount of walking performed to be reduced so markedly that the overall energy cost would be decreased. We controlled for any other factors that may cause changes in amount of walking, such as subjects’ schedules, means of transportation, and day of the week. The results of this study not only have implications for prescribing exercise regimens to improve public health and promote weight loss, but also help understand the expected impact of energy-saving prostheses and orthoses on exercise obtained during walking.

## METHODS

In this study, subjects wore two pairs of shoes, one pair weighted to increase the metabolic rate of walking by approximately 25% and one pair unweighted. The shoes were worn for one week each while walking activity was measured using shoe-mounted sensors. Functions relating metabolic rate to walking speed were calibrated to each subject with each pair of shoes in a laboratory setting and used in combination with walking activity data to calculate daily energy expenditure. A post-collection survey of participants’ subjective response to the shoes was also administered. Total metabolic energy expenditure, distance traveled, strides taken, average velocities, and survey answers were then compared.

### Shoe construction

Two pairs of flat-soled sneakers were constructed; one pair weighted and one pair unweighted. A high density foam platform (Iron Cloud EVA, Atlas International) was attached to the sole to house an inertial measurement unit (IMU; Sapphire Inertial Monitor, APDM, Inc.) (Fig. 1). The shoes were water resistant. A steel bar of appropriate mass was built into the weighted shoes, calibrated to equal approximately 2.5% of each user’s body mass, or 1.25% of body mass per foot. Mass ranged from 0.73 kg to 0.92 kg per foot, expected to correspond to a 25% increase in energy cost per step for each participant.^15^ Shoes were designed to avoid interfering with other normal foot functions.

**Figure 1:**
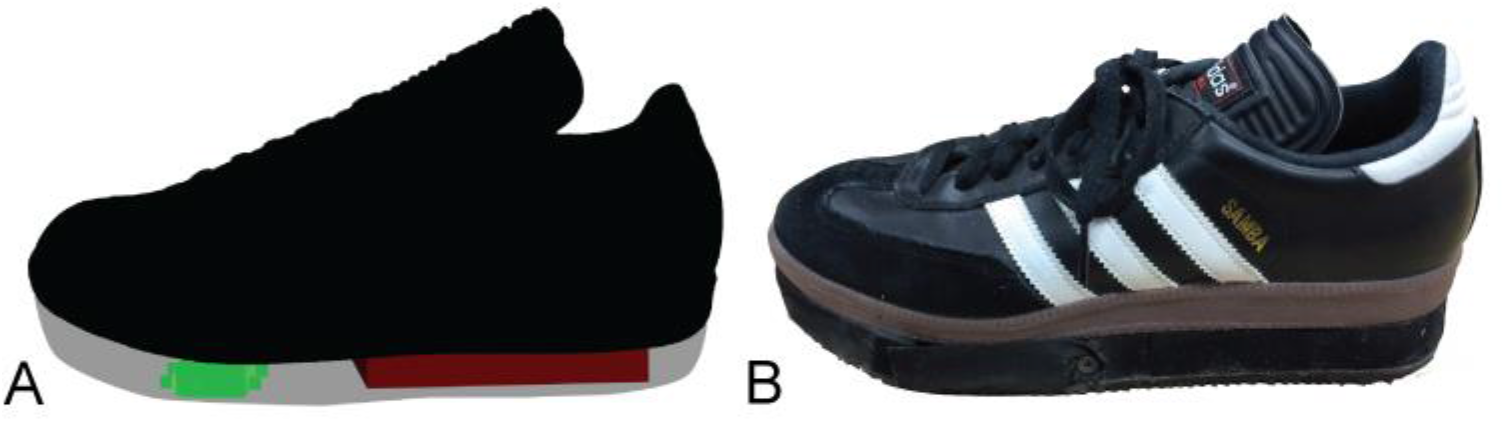
The Weighted Shoe. (**A**) Illustration of the weighted shoe, with an inertial measurement unit (light grey) and steel bar (dark grey) embedded in the sole. The mass of the steel bar was selected so as to increase energy cost of walking for the subject by 25%. (**B**) Photograph of one weighted shoe. An access panel in front of the inertial measurement unit allowed access for recharging.

### Participant information

The study protocol was approved by the Carnegie Mellon University Institutional Review Board, and written informed consent was obtained from all subjects after the nature and possible consequences of the study were explained. A power analysis of data from prior experiments suggested that about ten subjects would be required to discriminate differences of 10% in energy consumption with 50% power. Twenty-four participants with no cardiovascular, respiratory, metabolic or orthopedic conditions were recruited at Carnegie Mellon University. Five participants voluntarily withdrew before completion of the protocol, primarily citing dislike of the weighted shoes. An additional eight participants were excluded because they did not wear the shoes for the entire day (defined as at least 12 hours per day) or spent large periods of time with the shoes off during the day (defined as at least four hours), which indicated failure to comply with the study protocol and prevented measurement of all walking activity. One additional subject was excluded because environmental conditions changed between the two weeks (access to a car one week but not the other).

Data from the remaining subjects (N = 10, 7 male, 3 female, 23.6 ± 2.9 years, 22.9 ± 2.5 kg·m^-2^ body mass index) were included in the analysis. Height and weight were self-reported. Data from five complete, matched days of the week were used for each set of shoes. Matching days of the week, such as Monday in weighted shoes compared to Monday in unweighted shoes, was intended to control for the effects of daily schedules.

### Community-based collection

Participants were asked to wear the shoes all day, from the time they got out of bed until the time they returned, except when bathing. Activities that required the use of other shoes or no shoes, such as running or swimming for exercise, were logged and kept uniform between the two weeks to keep other sources of exercise-related energy expenditure consistent. Two subjects had additional activities to report, which were consistent between shoe conditions. Subjects were provided with a wall charger and asked to charge the IMU nightly. Participants were not informed of the exact motivations behind the study so as to avoid triggering conscious decisions to alter walking behavior.

Subjects were randomly given one pair of shoes for the first week and the other pair for the second week. Seven participants started with the weighted shoes and three with the unweighted shoes. We expected that behavioral changes would occur quickly, surfacing within the first week of exposure to the weighted shoes, and likewise would recede quickly, limiting interference effects.

Shoes were provided to subjects for one week each, but only the last five full, matched days were analyzed for each shoe, because five subjects only had five usable days. The other five subjects had six usable days, but the last five were used for consistency and to obtain the most acclimated of the possible five-day sets. Each set of shoes was worn for at least 12 hours in each day included in this analysis.

### Survey of subjective experience

After wearing both pairs of shoes, participants answered three pairs of qualitative questions for each pair:

1. How did the heavier (lighter) shoes affect your ease of walking? (Easier/Harder to walk)
2. The heavier (lighter) shoes made me feel more tired than usual. (Agree/Disagree)
3. If I were forced to do so, I would not mind having to wear the heavier (lighter) shoes for the rest of my life. (Agree/Disagree)

The survey utilized a nine point scale from -4 to 4 to determine participants’ opinions, where a value of 0 was neutral.

### Laboratory-based collection

At the conclusion of the community-based data collection, subjects walked on a treadmill while wearing each set of shoes at a range of speeds while oxygen consumption was measured. Subjects were instructed to fast for four hours prior to the collection. A three-minute quiet standing condition was collected to determine resting metabolic rate. Subjects then walked at three different speeds per shoe: 1.0, 1.25 and 1.5 m·s^-1^. The order in which the shoes were worn was randomized, as were the order of speeds for each shoe. Each condition lasted six minutes, with a two minute rest period between conditions. Whole body oxygen consumption was measured for each condition using an indirect respirometry system (Oxycon Mobile, CareFusion).

### Data analysis

Oxygen consumption and carbon dioxide production for each condition were averaged and entered into a standard formula to determine metabolic rate^16^. Net metabolic rate was calculated by subtracting the rate for quiet standing from each walking condition. The relationship between walking speed and net metabolic rate was then approximated for each subject using a linear model^17^, with zero net metabolic rate at zero velocity and a subject-specific slope that minimized the root-mean squared error from experimental data collected in the laboratory.

IMU data were visually examined to ensure that the shoes were worn all day, and that any quiet periods in the IMU corresponded to logged shoeless activities. Data was then numerically integrated using IMUWalk (Intelligent Prosthetic Systems, LLC) to estimate the length and duration of each walking stride, from right heel strike to right heel strike^18^. Measurements of vertical displacement, related to walking slope, were found to be unreliable during calibration tests and were not used. Walking velocities were calculated as stride length divided by stride period. Strides with velocity of less than 0.5 m·s^-1^ or greater than 2.5 m·s^-1^ or length of less than 0.15 m or greater than 2 m were excluded from analysis to ensure that all strides were due to walking, not foot tapping, shocks to the accelerometer or misclassification of activity. Energy cost per step was calculated by multiplying the speed-specific net metabolic rate for each step by the duration of that step.

We compared total energy cost, step count, average velocity, and average distance across shoe conditions, both for the entire five day period and for individual matched days. Paired t-tests with a significance level of α = 0.05 was used to determine statistical significance.

## RESULTS

The shoes were worn for 13.9 ± 1.0 hours per day (mean ± st. dev.) in the weighted shoe condition and 14.3 ± 1.3 hours per day in the unweighted shoe condition. The weighted shoes increased metabolic rate by 26.4±14.5% while walking at 1.25 m·s^-1^ (P = 3·10^−4^; paired t-test).

Survey results revealed a strong dislike for the weighted shoes (Fig. 2). All subjects disliked the weighted shoes compared to the unweighted shoes (1.8 ± 2.6 weighted versus -2.5 ± 1.3 unweighted, where 4 is strong dislike; P = 0.001). All subjects found it more difficult to walk with the weighted shoes, (2.6 ± 1.1 weighted versus -1.2 ± 1.8 unweighted, where 4 is much more difficult; P = 6·10^−4^). Most subjects found the weighted shoes to be more tiring than the unweighted shoes (2.1 ± 1.4 weighted vs. -1.7 ± 1.8 unweighted, where 4 is much more tiring; P = 0.003).

**Figure 2:**
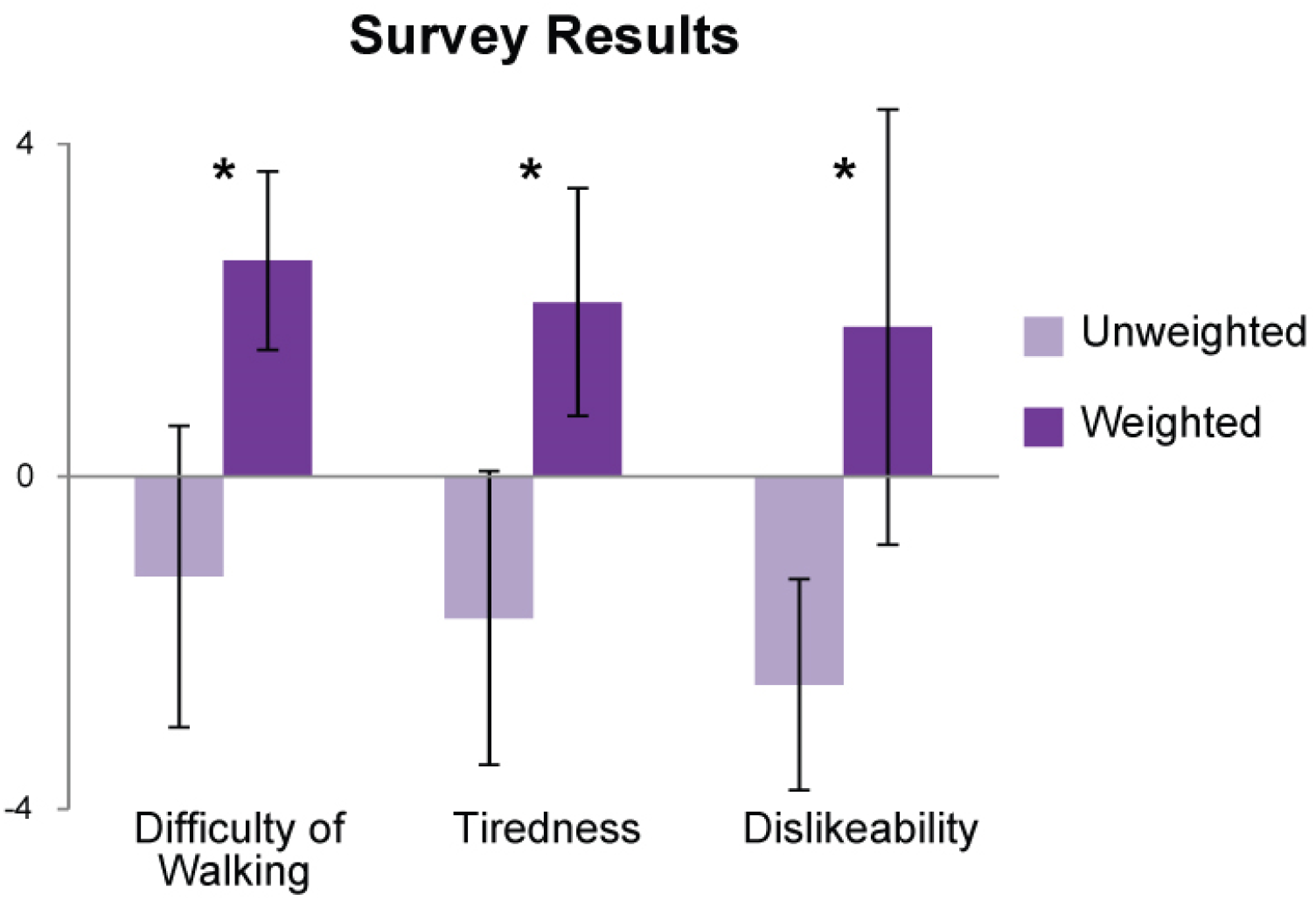
Survey results. All questions were based on a scale from -4 to +4, where +4 indicated extreme difficulty of walking, extreme tiredness, or extreme dislike of the shoes. Users uniformly disliked the weighted shoes and found them more difficult to walk in and more tiring (P ≤ 0.01). Light grey indicates data regarding the unweighted shoes. Dark grey indicates data regarding the weighted shoes.

Despite participants’ strong dislike of the weighted shoes, the average distance walked over five days was only reduced by 6%, and this was not a statistically significant change (2,700 ± 2,000 m·day^-1^ weighted vs. 2,900 ± 2,700 m·day^-1^ unweighted; P = 0.6; Fig. 3). The large standard deviations in these values indicate large inter-subject variability (Fig. 4). The number of steps per day was reduced by 4%, also not a statistically significant change (5,100 ± 2,500 steps·day^-1^ weighted vs. 5,300 ± 2,500 steps·day^-1^ unweighted; P = 0.7). There was no significant difference in average walking velocity (1.07 ± 0.14 m·s^-1^ weighted vs. 1.07 ± 0.17 m·s^-1^ unweighted; P = 0.9). The number of steps taken per day by our participants was similar to the national average of 5,800 steps taken by 18-29 year olds.^19^

**Figure 3:**
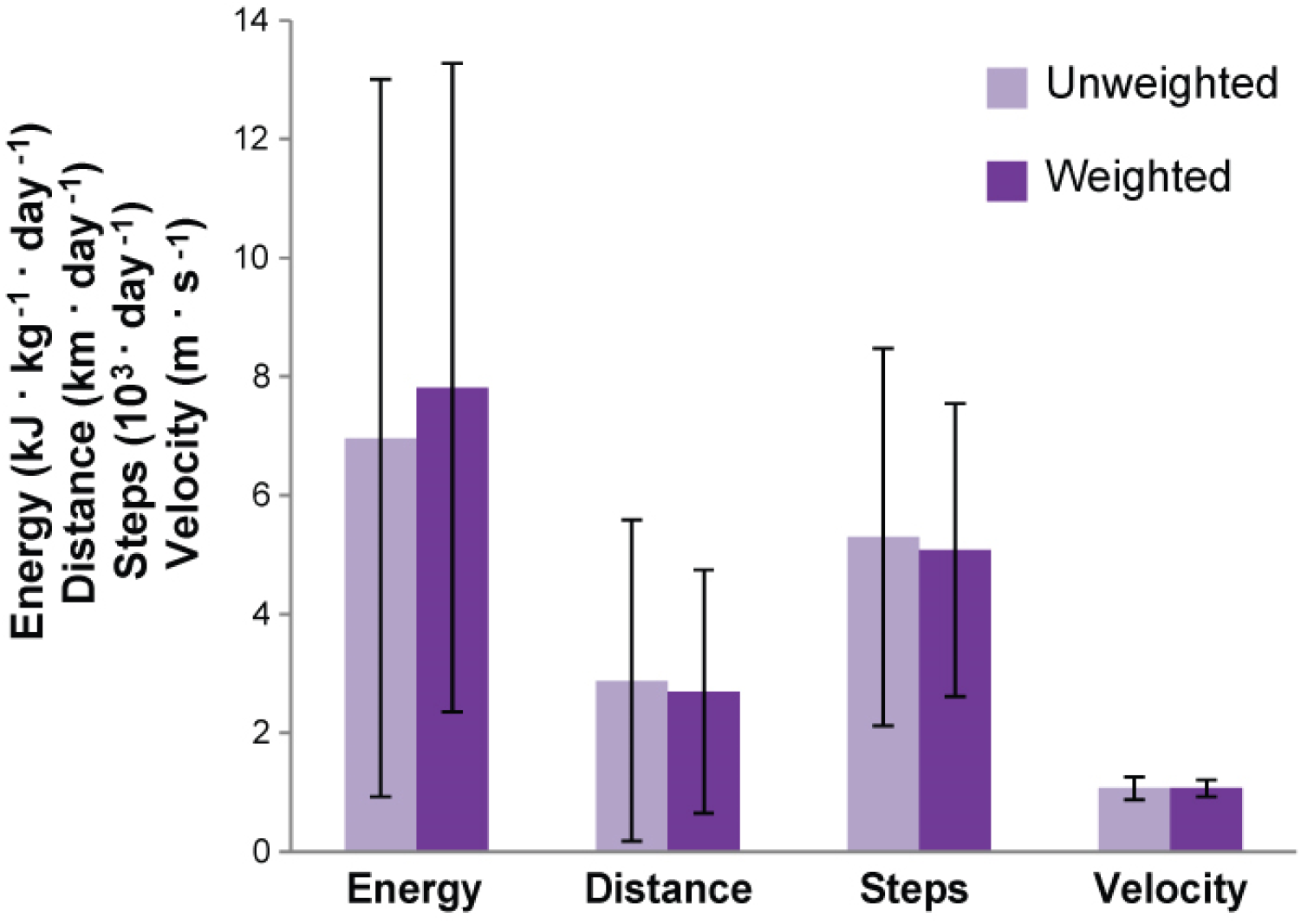
Average Values. Comparison of walking activity during the week-long community-based data collection. Metabolic energy expended through walking each day appeared to increase with the weighted shoes (P = 0.2). This result was driven by the increase in energy cost per step, coupled with only small decreases in distance walked, steps per day and average speed, none of which were statistically significant (P ≥ 0.6). Light grey indicates data regarding the unweighted shoes. Dark grey indicates data regarding the weighted shoes.

**Figure 4:**
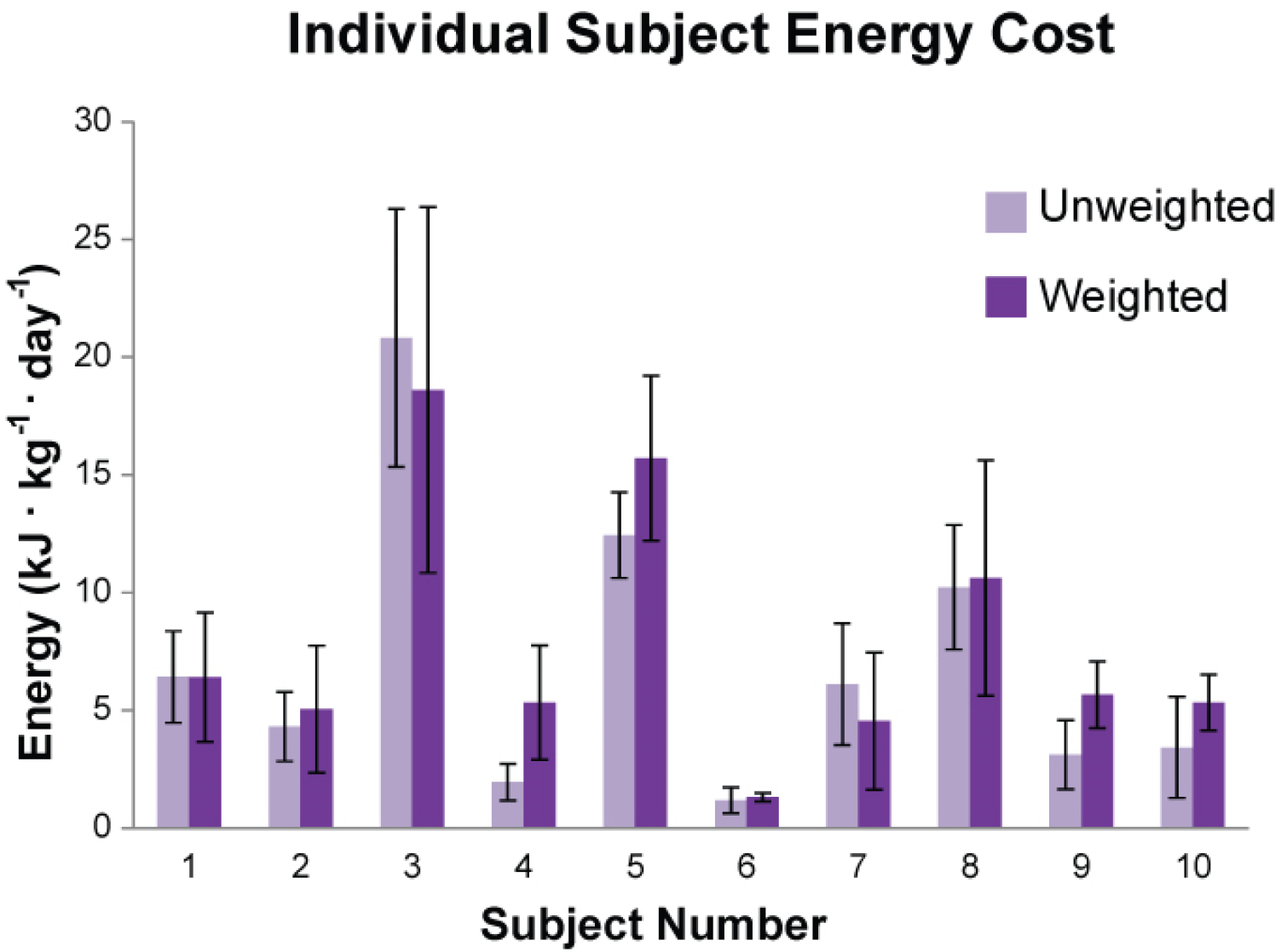
Individual Energy Costs. Subject-wise comparison of metabolic energy expenditure from walking during the week-long community-based data collection. Energy expenditure varied widely across subjects, but increased with the weighted shoes in all but two cases. Light grey indicates data regarding the unweighted shoes. Dark grey indicates data regarding the weighted shoes.

With substantially increased energy cost per step and smaller reductions in daily walking distance and velocity, the weighted shoes led to a 12% increase in energy expenditure over the five day collection period, although this result was not statistically significant (7,800 ± 5,500 J·kg^-1^·day^-1^ weighted vs. 7,000 ± 6,000 J·kg^-1^·day^-1^ unweighted; P = 0.2). All but two subjects used more energy with the weighted shoes than with the unweighted shoes (Fig. 4). Large standard deviations in these data are again due to inter-subject variability.

By the final day of the community-based collection, however, energy expended using the weighted shoes was 24% lower than with the unweighted shoes, although this difference was not statistically significant (6,400 ± 5,000 J·kg^-1^·day^-1^ weighted vs. 8,400 ± 6,400 J·kg^-1^·day^-1^ unweighted; P = 0.4). Distance walked (P = 0.3), steps taken (P = 0.2) and velocity (P = 0.7) also appeared to be reduced for the weighted shoes by the fifth day, but not significantly (Fig. 5).

**Figure 5:**
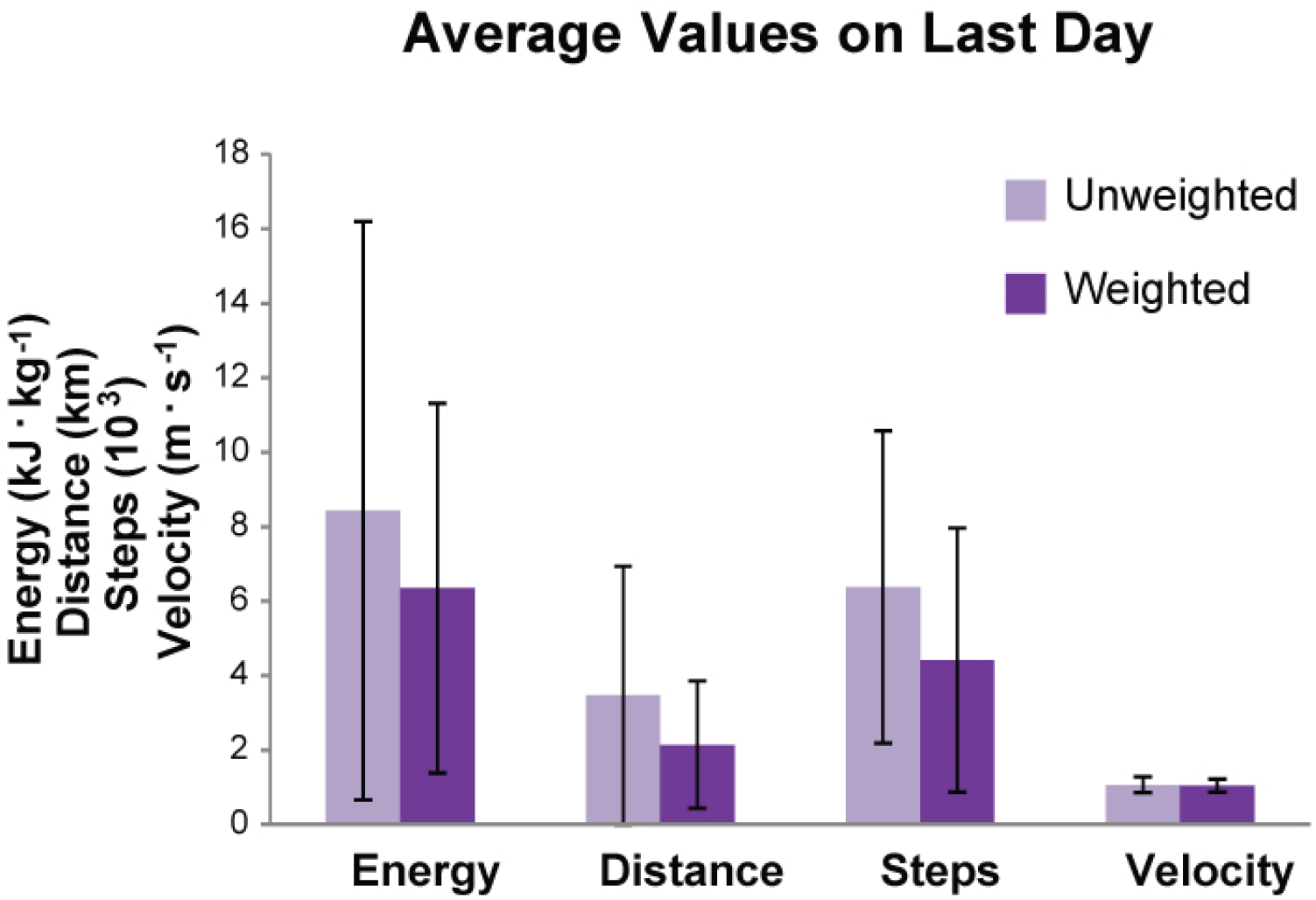
Average Values on Final Day. Comparison of walking activity during the final day of the community-based data collection. Metabolic energy expended through walking appeared to decrease with the weighted shoes (P = 0.4), driven by substantial, though not statistically significant, decreases in distance walked (P = 0.3) and number of steps (P = 0.2). Light grey indicates data regarding the unweighted shoes. Dark grey indicates data regarding the weighted shoes.

## DISCUSSION

The aim of this study was to examine the trade-off between the effort of walking on each step and the daily energy expenditure due to walking. We found a small, though not statistically significant, increase in energy expenditure over a five-day period when subjects wore weighted shoes that substantially increased the energy expended on each step. The implications of this finding for weight loss interventions are limited, however, because subjects uniformly reported a strong dislike of the weighted shoe condition, and many individuals recruited into the study dropped out or failed to adhere as a result. It would therefore likely be difficult to realize this small increase in exercise from weighted shoes under natural conditions. The results of this study did not support our hypothesis that behavioral changes to reduce walking activity would outweigh the effects of increased energy cost per step, but this finding is muted by experimental limitations. The most prominent practical limitations are the duration of the community-based data collection and self-selection of participants through dropout.

The community-based data collection in this study may not have lasted long enough to capture changes in self-selected walking activity in response to added energy cost per step. Humans tend to quickly select coordination patterns that minimize energy expenditure during walking,^20^ likely a result of evolutionary pressures opposing waste of energy.^21^ It is not yet understood whether similar pressures affect longer-term behavior, but there is some evidence that they may. For example, sedentary adults are more likely to adhere to a moderate-intensity walking regimen for six months than to a more vigorous routine^8^. As another example, people tend to shift the balance of their commuting activities towards mechanized transport if placed in a built environment that makes walking more difficult.^22^ It is not clear how long one must be exposed to an increase in energy cost before such changes take effect, however, and our results suggest that it requires more than one week. Several subjects in this study described beginning to change their schedules in ways that reduced the amount they walked while wearing the weighted shoes. For example, some took the bus or drove instead of walking, chose to stay at home instead of engaging in social events, or chose to miss the bus rather than run to catch it. This may help explain findings from the final day of the collection, in which subjects walked so much less with the weighted shoes that they expended less energy than with the unweighted shoes.

Participants included in this analysis may not represent a random sample of the population, but rather a subset that responded more favorably to the weighted shoe condition. About 60% of participants enrolled in this study either dropped out or were excluded due to a lack of adherence, primarily in the weighted shoe condition. Of the remaining individuals, some may have found the weighted shoes to be less objectionable, possibly because they were less influenced by energy cost. Others among the remaining individuals may have had psychological traits that made them more determined to complete the study, which may also have led them to continue their daily regimens despite the added effort. Correspondingly, recruits who dropped out may have been so strongly influenced by energy cost that they made the behavioral change of leaving the study to avoid it. Such self-selection may have led to a bias towards increased daily energy expenditure with the weighted shoes among the participants included as compared to the population as a whole.

These outcomes might also be influenced by the demographic traits of participants. Young, fit undergraduate and graduate university students may respond differently to changes in walking effort than the older, sedentary adults for whom obesity rates are highest.^1^ For example, it may be that more fit individuals are more tolerant of large increases in energy expenditure, or that individuals with fixed routines due to class schedules have less flexibility in selecting the amount of walking they perform. One would again expect such differences to result in more energy expended with the weighted shoes for the population tested compared to a broader population of adults.

The sample size obtained for analysis in this study was small given the large amount of inter-subject variability observed. Based on prior laboratory-based collections, we expected to detect moderate changes in metabolic rate with acceptable power using data from ten subjects. Inter-subject variations in daily walking activity were much larger than inter-subject differences in metabolic rate on each step, however, leading to large variability. The use of repeated measures or paired statistical tests mitigates but does not eliminate this issue. For example some subjects walked ten times as much as others over the course of a week, which allows mean differences in energy cost to differ substantially from differences in mean energy cost.

Future studies could attempt to address these issues using longer data collections, applying greater social or economic incentives to increase retention, and recruiting a larger number of obese adults. Of course, each of these improvements would be challenging for practical reasons. For example, we found that longer community-based data collection periods resulted in even greater dropout rates. We tried several different ways of incentivizing subjects over a four-year period prior to the beginning of the study, but were unable to obtain better retention. We were limited in the number of subjects we could recruit and test because of the expense of, e.g., fabricating custom shoes for each participant and carefully analyzing hundreds of hours of data for each one.

There were also technical limitations due to the hardware used to track subjects’ walking activity in the community-based portion of the experiment. The IMU and data logger we used needed nightly charging to ensure adequate battery life, and most subjects failed to properly charge the shoes on one or more nights, leading to a loss of data. Calibration tests with the IMU revealed that it did not track motion in the vertical direction with sufficient accuracy, so energy expenditure due to walking up stairs or slopes was not captured. Future experiments could use a lower-power data logger, a more accurate IMU, and GPS to avoid data loss and better estimate altitude changes on each step.

While the present results do not support the use of weighted shoes for increasing exercise obtained from walking, the instructions given to subjects may have an important effect. Adherence to a lifestyle change is heavily influenced by readiness and desire to change.^5,7,11^ Our subjects were not told the motivations behind the study in order to prevent intentional behavioral changes. It is possible that, had they known the study’s background, those who wanted to lose weight or increase physical activity would have intentionally walked the same amount or more with the weighted shoes to increase total energy expenditure.

This study may also have implications for assistive device design. Research on lower-limb prostheses and exoskeletons has recently focused on reducing the metabolic energy expenditure of walking.^23,24^ However, many prosthesis users are affected by obesity-related comorbidities, and 70% of all lower-limb amputations are due to complications from dysvascular diseases such as diabetes mellitus.^25^ Devices that reduce the metabolic energy expended on each step could reduce exercise from walking, worsening such conditions over the long term, if the amount of walking remained constant. There are conflicting indications as to whether prosthesis function affects amount of walking. For example, more comfortable socket liners lead to increased physical activity,^26^ while preferences across ankle-foot prostheses do not correspond to level of physical activity.^27^ The present results suggest that increased cost per step might increase exercise from walking over short periods, but might reduce the amount of walking over longer periods. A similar study conducted among amputees for a longer duration, and addressing the other limitations discussed above, would be helpful in understanding the long-term health effects of exoskeletons and prostheses that reduce the energy cost of walking.

## CONCLUSIONS

Walking constitutes a significant portion of the daily energy expenditure of humans and increasing the metabolic energy used during walking has the potential to help address obesity. In this study, we found that substantially increasing energy cost per step led to a smaller increase in the energy expended through walking over a five day period, partially offset by decreases in the amount of walking performed. Given the strong dislike of increased walking effort, however, it is unlikely that people would adopt such devices in daily life unless they were given strong external motivations to use them. Increases in overall energy use might be reduced or even reversed over time due to changes in self-selected walking activity, although the one-week duration of this study appears to have been insufficient to capture such changes. Further study of the long-term behavioral effects of increasing the energy cost of walking are merited, as they may lead to new interventions for obesity and would help us to understand the expected health effects of assistive devices that make walking easier.

## ACKNOWLEDGMENTS

This material is based upon work supported by the National Science Foundation under Grant No. IIS-1355716. The authors thank Josh Caputo, Ruthika, Solomon Abiola, Felix Chiu, and Shweta Singh for assistance with hardware and software development.

